# Proteomic Profiling of Hypoplastic Lungs Suggests an Underlying Inflammatory Response in the Pathogenesis of Abnormal Lung Development in Congenital Diaphragmatic Hernia

**DOI:** 10.1101/2022.03.07.483298

**Authors:** Richard Wagner, Paula Lieckfeldt, Hadeesha Piyadasa, Moritz Markel, Jan Riedel, Camelia Stefanovici, Nicole Peukert, Daywin Patel, Gabrielle Derraugh, Suyin Lum Min, Jan-Hendrik Gosemann, Jan Deprest, Christopher D. Pascoe, Andrew Tse, Martin Lacher, Neeloffer Mookherjee, Richard Keijzer

**Affiliations:** Departments of Surgery, Division of Pediatric Surgery, Pediatrics & Child Health, University of Manitoba, Winnipeg, Manitoba, Canada; Department of Physiology and Pathophysiology, University of Manitoba, Winnipeg, MB, Canada; Biology of Breathing Theme, Children’s Hospital Research Institute of Manitoba, Winnipeg, Manitoba, Canada; Department of Pediatric Surgery, University Hospital Leipzig, Leipzig, Germany; Manitoba Center for Proteomics and Systems Biology, Department of Internal Medicine, University of Manitoba, Winnipeg, Manitoba, Canada; Department of Immunology, University of Manitoba, Winnipeg, Manitoba, Canada; Department of Pathology, University of Manitoba, and Shared Services Manitoba, Winnipeg, MB, Canada; University Hospitals KU Leuven, and Department of Development and Regeneration, Biomedical Sciences, KU Leuven

**Keywords:** Congenital Diaphragmatic Hernia, Lung Development, Proteomics, Inflammation, Nitrofen Rat Model, FETO

## Abstract

The pathogenesis of lung hypoplasia in congenital diaphragmatic hernia (CDH), a common birth defect, is poorly understood. The diaphragmatic defect can be repaired surgically, but the abnormal lung development contributes to a high mortality in these patients. To better understand the underlying pathobiology, we used the nitrofen rat model of CDH and characterized the proteome of hypoplastic CDH lungs at the alveolar stage (E21). Amongst the 218 significantly altered proteins between CDH and control lungs were Tenascin C, CREBBP, LYN and STAT3. We showed that Tenascin C was decreased around the distal airway branches in nitrofen and human fetal CDH lungs. In contrast, STAT3 was significantly increased in the airway epithelium of nitrofen lungs at E21. STAT3 inhibition after direct nitrofen exposure to fetal rat lung explants (E14.5) partially reversed the hypoplastic lung phenotype *ex vivo* by increasing peripheral lung budding. Moreover, we demonstrated that several STAT3 associated cytokines (IL-15, IL-9, IL-2) are increased in fetal tracheal aspirates of CDH survivors compared to non-survivors after fetoscopic tracheal occlusion. Using pathway analysis for significantly altered proteins in our proteomic analysis, we observed an enrichment in inflammatory response associated with Epstein Barr Virus and cytokine signaling in nitrofen CDH lungs. However, we were unable to detect EBV mRNA via *in-situ* Hybridization in human CDH lungs. With our unbiased proteomics approach, we show for the first time that inflammatory processes are likely underlying the pathogenesis of abnormal lung development in CDH.

## Introduction

Congenital Diaphragmatic Hernia (CDH) is a severe birth defect that is characterized by a hole in the diaphragm that manifests during fetal development. Lung hypoplasia in CDH is associated with reduced airway branching and abnormal alveolar and mesenchymal architecture ^1^. Despite improved postnatal medical and surgical care, mortality due to lung hypoplasia and persistent pulmonary hypertension remains high (30% - 50%) ^2^. With an incidence similar to cystic fibrosis (1:2500), CDH accounts for 8% of congenital anomalies, but in contrast to cystic fibrosis the underlying pathogenesis of CDH is unclear. A monogenetic cause has not been identified for CDH, suggesting that epigenetic and environmental factors are involved ^3^.

Administration of the herbicide nitrofen to pregnant rats causes rendition of the CDH phenotype (lung hypoplasia + diaphragmatic hernia) in 50% of the offspring, but it remains unclear how nitrofen works in this model ^4^. In humans there is no correlation between nitrofen exposure and CDH. In rat and human studies, we have previously demonstrated that an increase in microRNA 200b – a post-transcriptional regulator - is associated with better outcomes and improved lung maturation in CDH cases ^5–7^. Others have shown that different genes are altered at transcriptional and posttranslational levels in hypoplastic CDH lungs, but the used methods were predominantly single-plexed and antibody-based ^8^. We and others have used early “omics” - approaches to determine the transcriptome of nitrofen-induced hypoplastic lungs ^9,10^. Yet, these studies were unable to explain the pathogenesis of CDH. Shotgun-proteomics allow for simultaneous analysis of thousands of proteins in a multiplexed and unbiased manner with increasing throughput and accuracy ^11,12^.

Here, we used a proteomics approach with LC-MS/MS to determine changes in the lung proteome of nitrofen-induced hypoplastic CDH lungs compared to controls. Using pathway analysis, we observed an enrichment in signalling pathways associated to inflammatory processes (e.g., JAK/STAT, NFκB) and cytokine signalling within CDH lungs. Transcription factor STAT3 was a central hub of the interaction network of proteins that were altered, and was significantly increased in nitrofen CDH lungs. Direct STAT3 inhibition in *ex vivo* lung explants increased the total number of distal lung buds in nitrofen-induced lung hypoplasia. Moreover, we demonstrated that different cytokines that are associated with STAT3 signaling (e.g., IL-9, IL-2, IL-15) were significantly increased in fetal tracheal aspirates of patients surviving CDH compared to non-survivors.

Our study provides a comprehensive analysis of changes in the protein profile of hypoplastic CDH lungs and shows that key immune signaling intermediates and associated inflammatory cytokines are activated in abnormal lung development associated with CDH. Thereby, we propose a novel disease mechanism theory for the pathogenesis of lung hypoplasia in CDH.

## 2. Methods

### 2.1 Ethical approval and sample acquisition

This study was approved by the health research ethics board from the University of Manitoba and University of Leipzig (19-010/1/2 (AC11436) and HS15293 (H2012:134); T03/21-MEZ). Animal studies were compliant with the ARRIVE guidelines ^13^. Female Sprague Dawley rats were housed in the central animal care facility at the University of Manitoba. The rats were randomly distributed to individual cages by the animal care staff (1 rat per cage). Lung hypoplasia and CDH were induced using nitrofen as previously described ^4^. Briefly, pregnant dams were gavaged in the morning of embryonic day 9 (E9) with nitrofen + olive oil (CDH, n=5) or olive oil only (control, n=5). After 21 days of pregnancy (E21) rats were euthanized, fetal lungs were harvested and either stored at −80 °C or formalin fixed and paraffin embedded after 48h. Human paraffin embedded lung tissues from CDH patients and controls were obtained from a previously established biobank at the University of Manitoba ^14^. For lung explant culture, fetal lungs were excised one-by-one at E14.5 and cultured for 2-3 days. The lung explants were randomly assigned to either DMSO, nitrofen or nitrofen + STAT3 inhibitor treatment. We previously determined that DMSO, which served as vehicle for nitrofen, had no adverse effects compared to control medium (Suppl. Figure 1) ^15^. Tracheal aspirate samples from FETO CDH cases were obtained from 8 patients (4 that subsequently survived CDH and 4 non-survivors) at 28 weeks’ gestation as described before ^5^ (Suppl. Table 1).

### 2.2 Sample processing and LC-MS/MS

Samples were processed for LC-MS/MS as previously described ^16^. Briefly, flash frozen lung samples were homogenized in protein extraction buffer containing 1% SDS and protease inhibitor cocktail (Cell signalling) and centrifuged at 10,000G. Lysates were collected, and aliquoted and protein concentration was determined using the Micro BCA Protein Assay (Thermo Fisher Canada). Single-pot-solid-phase-enhanced-sample preparation (SP3) was used for protein clean up. 100 μg tissue lysates were reduced with 10mM DTT (final concentration) at 57°C for 30 minutes and alkylated with 50mM IAA (final concentration) for 45 min at RT. Samples were digested with trypsin (Promega) at an enzyme to protein ratio of 1:25 (w/w) for 16 hours at 37°C. The resultant peptide mixture was desalted using a SOLA™ HRP SPE cartridge (Thermo Fisher) and lyophilized. The final peptide concentration was determined using NanoDrop 2000 (Thermo Fisher). Peptide separation was performed on a C18 (Luna C18(2), 3 μm particle size (Phenomenex, Torrance, CA)) column packed in-house in Pico-Frit (100 μm X 30 cm) capillaries (New Objective, Woburn, MA). Analyses of peptide digests were carried out on an Orbitrap Q Exactive HF-X instrument (Thermo Fisher Scientific, Bremen, Germany). The instrument was configured for data-dependent methods using the full MS/DD?MS/MS setup in a positive mode. Survey scans covering the mass range of 375–1500 m/z were acquired at a resolution of 120,000 (at m/z 200).

Peptides were identified and quantified with X!Tandem (cyclone 2012.10.01.1) using the following settings: Variable modification M, W + 15.995 (oxidation) or +31.989 (Double oxidation); constant modification C+57.021 (Cystine protection); S, T, Y +79.966 (phosphorylation); N, Q +0.984 (deamidation); fragment mass error 0.1 DA; parent mass error ±20 PPM. We selected peptides with a minimum of two unique peptide sequences and confidence values of log(e)<−3. For label-free quantitation of every selected protein, the sum of the MS/MS fragment intensities of each associate peptide were used and transformed into a log2 scale for subsequent differential expression analysis.

### 2.3 Western Blot

Equal amounts (10 μg, *ex vivo:* 20 μg) of protein lysates as described above were resolved on 4- 12% Bis-Tris Protein gels (Invitrogen). Proteins were transferred to nitrocellulose membranes (Millipore, Billerica, MA) or PVDF membranes (*ex vivo;* BIO-RAD), blocked with 5% (w/v) milk powder dissolved in TBST (20mM Tris–HCl pH 7.5, 150mM NaCl, 0.1% Tween-20) and probed with antibodies against LYN (Abcam), Tenascin C (Cell Signalling) and STAT3 and pSTAT3(Tyr705) (Cell Signalling) in 1% (w/v) milk powder (Lyn, Tenascin C) or 5% BSA (pSTAT3, STAT3; Serva) dissolved in TBST. Affinity purified horseradish peroxidase-linked secondary antibodies (Cell Signaling) were used for detection. Membranes were developed with Amersham ECL/Prime or Select detection systems (GE Healthcare) according to the manufacturer’s instructions. Band intensities were normalized against Glyceraldehyde-3-phosphate dehydrogenase (GAPDH).

### 2.4 Immunofluorescence

Immunofluorescence (IF) on rat and human lung tissues were carried out on 6 μM sections of formalin fixed tissues as previously described ^5,7^. Primary antibodies against Tenascin C (Cell Signalling) CREBBP (Cell Signalling) and STAT3 (Cell Signalling) were used for IF.

### 2.5 In-Situ-Hybridization

Human tissue sections were subjected to an automated stainer (Benchmark XT, Ventana Medical Systems) using an INFORM EBER probe (Ventana Medical Systems, Tucson AZ, USA) as previously described ^17^. After ISH for detection of Epstein Barr Virus (EBV), slides were examined by two independent pathologists who were blinded to the disease status of the samples.

### 2.6 Retrospective case-control study

We performed a retrospective case-control study of women who gave birth to a baby with Bochdalek-type congenital diaphragmatic hernia between 1991 and 2015. We created a link between the Manitoba Centre for Health Policy (MCHP) repository and the Winnipeg’s Surgical Database of Outcomes and Management (WiSDOM) to establish the cohort. The control group was created using a 1:10 date-of-birth matched population from the MCHP database. An ICD-9-CM code of 075 and 078 were used for the diagnosis of infectious mononucleosis and cytomegaloviral disease, respectively. The linkage was done using SAS^®^ software and the statistical analysis was conducted using R^®^ software. A Fischer’s exact test and p<0.05 was used to determine statistical significance.

### 2.7 Cytokine Array

Tracheal aspirate samples from 8 CDH patients (4 survivors vs. 4 non-survivors) undergoing FETO were profiled using the 42-plex Discovery assay (Human Cytokine Array/Chemokine Array 42-Plex Panel, Eve Technologies, Alberta, Canada).

### 2.8 Bioinformatic Analysis and Statistical Analysis

Volcano plot, PLS-DA and VIP-score/ Heat map were generated using the R-Software package (ggplot) and correlation analysis was performed with the ggully, ggpubr and ggally package ^18,19^. T-test, Mann-Withney test or ANOVA test were performed when indicated. Statistical significance was defined as p<0.05. A total of 218 proteins that were changed between CDH and control lungs with a p-value <0.05 and log fold change > 1.5 were assessed with pathway analysis via NetworkAnalyst 3.0 ^20^ and Ingenuity Pathway Analysis (IPA) tool (Qiagen Inc) ^21^.

## 3. Results

### 3.1 Nitrofen CDH lungs exhibit an altered protein profile during the alveolar phase of lung development

The well-established nitrofen rat model mimics the human CDH phenotype (lung hypoplasia + diaphragmatic hernia), but the molecular mechanisms in this model remain enigmatic ^22^. To better understand underlying pathobiological processes, we compared global protein profiles of nitrofen-induced hypoplastic CDH lungs (n=5) and controls (n=5) at the alveolar stage (E21) of lung development (Figure 1A). Using label-free 1D LC-MS/MS we identified 4046 proteins with high confidence (proteins with a confidence value of log(e) <−3 with a minimum of two unique peptides). Of these, 218 proteins were significantly different, with 94 increased and 124 decreased proteins (by log2 >1.5, p<0.05) in hypoplastic CDH compared to normal lungs (Figure 1B). Partial least squares discriminant analysis (PLS-DA) separated the dataset into 2 independent groups based on their lung proteomic profiles (Figure 1C). We followed this with a VIP-score (Variable importance of projection) analysis to determine the proteins that contributed most significantly to the segregation seen in our PLS-DA analysis. This revealed 1096 proteins with a VIP-score > 1, and 93 proteins with VIP-score > 2. The VIP-scores of the top 20 of our altered proteins together with a corresponding heat map are displayed in Figure 1D. From the top 20 proteins, only 2 proteins (Tenascin C, Keratin 4) were decreased in hypoplastic nitrofen lungs. Tenascin C was significantly suppressed in nitrofen lungs and had the highest VIP-score (VIP-score = 2.8; logFC = −1.73; p < 0.0001), whereas CREB-binding protein (CREBBP) showed enhanced abundance (VIP-score = 2.1; logFC = 2.1; p < 0.001). Creating a minimum protein interaction network with an input of all 218 significantly changed proteins, we found that CREB-binding protein (CREBBP), Tyrosine-protein kinase LYN (LYN), Signal transducer and activator of transcription 3 (STAT3) and nuclear factor kappa-light-chain-enhancer of activated B cells 1 (NF-kB1) were central protein hubs in this network (Figure 1E).

**Figure 1:**
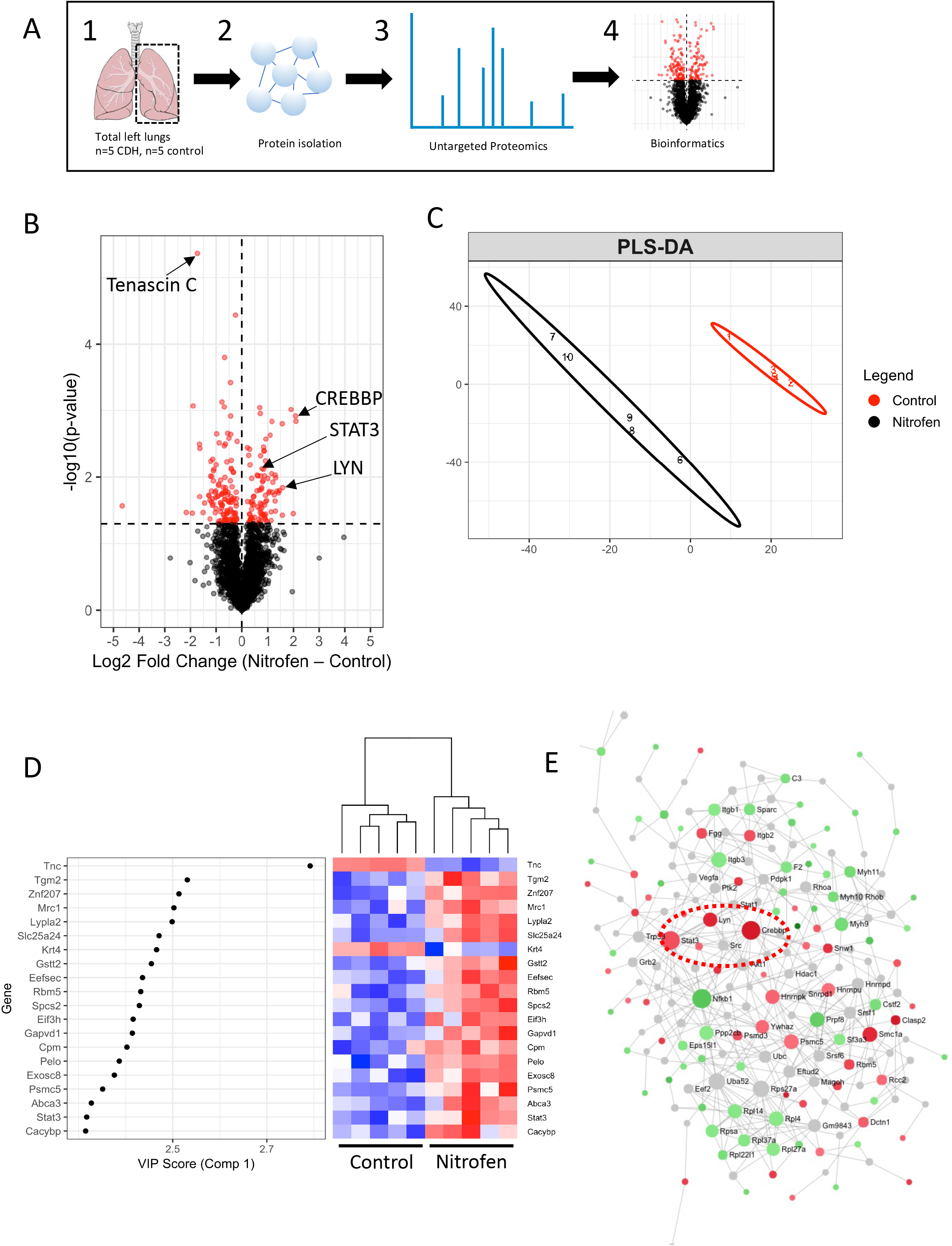
Nitrofen CDH lungs exhibit an altered protein profile during the alveolar phase of lung development. A) Schematic of the work flow: In vivo nitrofen exposed left lungs with full CDH phenotype (n=5) and control lungs (n=5) were obtained at E21, total proteins were isolated followed by untargeted proteomics and bioinformatic analysis B) Volcano plot highlighting 218 proteins that were significantly altered (red) between nitrofen and control lungs, 94 increased and 124 decreased (log2FC>1.5; p< 0.05). C) Partial least squares discriminant analysis (PLS-DA) of the proteomic dataset D) VIP-scores (Variable importance of projection) with correlating heat map showing the protein targets with the strongest contribution to the separation of the two groups determined by PLS-DA E) Protein interaction network with all significantly altered proteins between nitrofen CDH and control lungs created via NetworkAnalyst3.0. Highlighted are CREBBP, LYN and STAT3 as central protein hubs within this network.

### 3.2. Independent confirmation of selected proteins demonstrates decreased Tenascin C in nitrofen and human CDH, and enhanced abundance of CREBBP, LYN and STAT3 in nitrofen CDH lungs

In line with the proteomic analysis, immunostaining against Tenascin C showed a decrease in nitrofen lungs, especially around the distal airway branches, where it was highly expressed in control rat lungs (Figure 2A, n=3 per group). We confirmed decreased total protein abundance of Tenascin C in nitrofen lungs with Western blot (Figure 2B). To assess these observations in human CDH lungs, we performed Tenascin C immunostaining in fetal lung sections from pathology specimens and observed an analogous decrease in CDH lungs when compared to age matched controls (n=3 per group, Figure 2C).

**Figure 2:**
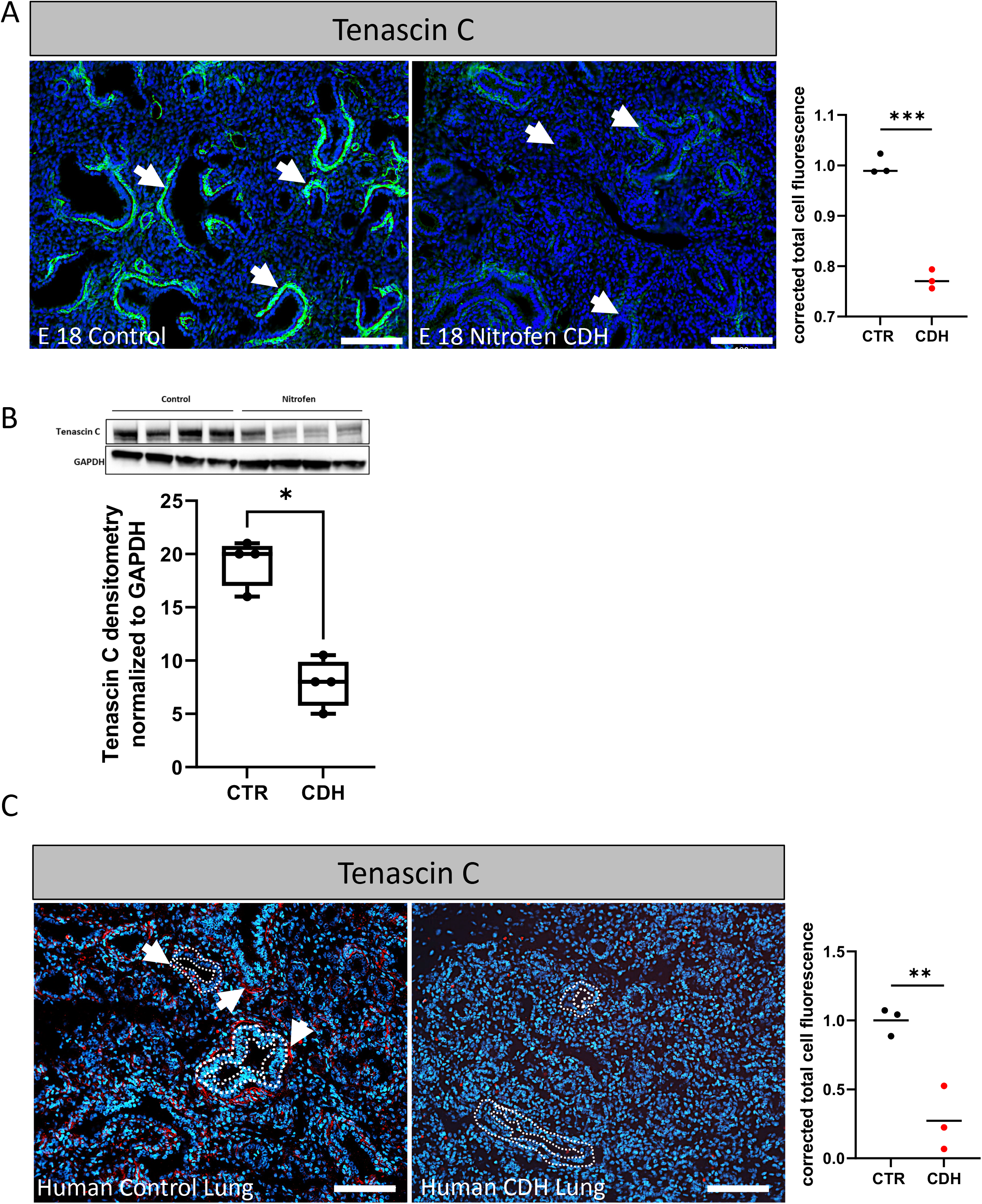
Tenascin C is significantly decreased in nitrofen and human fetal CDH lungs. A) Representative images for immunostaining against Tenascin C in nitrofen and control lungs at E18 (n=3 per group). Arrows indicating areas around the distal airway branches. Quantification via corrected total cell fluorescence B) Western blot with antibody against Tenascin C in E21 nitrofen and control lungs and quantification via densitometry (n=4 per group). C) Immunostaining against Tenascin C in human full-term CDH lungs (n=3) and controls (n=3) from fetal pathology specimens. Some distal airway branches are outlined. Quantification via corrected total cell fluorescence. Mann-Withney U test was used for statistical analysis, * p<0.05, ** p<0.01, *** p<0.001. Scale bars 100μm.

We further selected proteins that were identified as central hubs (CREBBP and LYN) in the interaction network of proteins that were significantly changed between nitrofen CDH and control lungs for independent validation studies using various immunoassays. Immunostaining with an antibody against CREBBP showed ubiquitously increased abundance of this protein in nitrofen lungs (Figure 3A). Moreover, Western blot for LYN confirmed the overall increase in nitrofen lungs (Figure 3B). Additionally, we generated a protein interaction network with the VIP-score Top 50 discriminant proteins between nitrofen and control lungs, and found that STAT3, which was increased in CDH, was the central node in this network (Figure 3C). Using immunostaining against STAT3, we detected enhanced abundance of STAT3 in the airway epithelium of nitrofen CDH lungs at E21 (Figure 3D).

**Figure 3:**
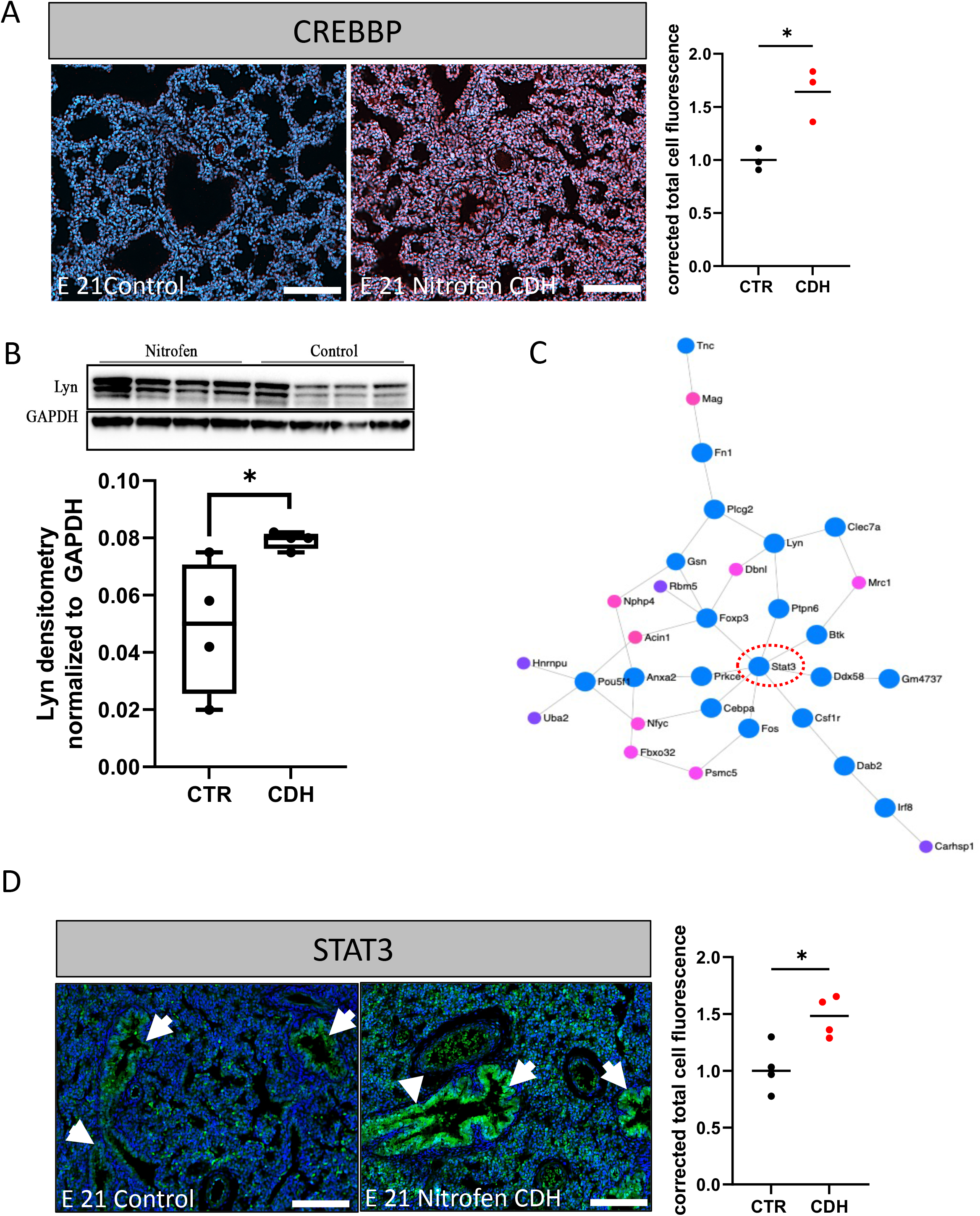
CREBBP, LYN and STAT3 are significantly enhanced in nitrofen CDH. A) Representative images of immunostaining against CREBBP in nitrofen CDH lungs at E21 (n=3 per group). Quantification via corrected total cell fluorescence. B) Western blot with antibody against LYN comparing E21 nitrofen and control lungs (n=4 per group). C) Protein interaction network using top 50 VIP discriminant proteins from the proteomics dataset. Proteins associated with immune response pathways are highlighted in blue. STAT3 highlighted as central protein hub for this network. D) Representative images of immunostaining against STAT3 in E21 nitrofen and control lungs (n=3 per group), Arrows indicating the respiratory airway epithelium. Mann-Withney U test was used for statistical analysis, * p<0.05. Scale bars 100μm.

### 3.3. Pathway analysis determines enrichment in processes associated to inflammation and immune response elements in CDH lungs

To assess pathways and biological processes that correspond to the 218 significantly altered proteins (94 decreased, 124 enhanced) in CDH lungs, we used NetworkAnalyst 3.0 and Ingenuity Pathway Analysis (IPA) bioinformatics tools. We found that the predominant pathways enriched were those associated with immune activation and known lung development pathways (e.g., Wnt, Notch; Figure 4A). Moreover, Kyoto Encyclopedia of Genes and Genomes (KEGG) pathway analysis highlighted pathways associated with EBV infection (Figure 4A). CREBBP, LYN, STAT3 and NF-κB were central elements within the EBV specific network of altered proteins in CDH lungs (Figure 4B). We thus hypothesized that prenatal nitrofen exposure triggers inflammatory pathways in developing rat lungs, similar to a viral stimulus to the innate immune system in human fetal CDH lungs. To test this hypothesis, we studied CDH and control fetopsy lung specimens for the presence of EBV mRNA using *in-situ* hybridization against the EBV encoding region (EBER). Two independent and blinded pathologists examined the stained sections and did not detect EBV mRNA in CDH or control lungs (Figure 4C). To further investigate potential correlations between EBV exposure during pregnancy and CDH occurrence, we performed a retrospective case-control study of women who gave birth to a baby with Bochdalek type CDH between 1991 and 2015. We used the Manitoba Centre for Health Policy and compared diagnosis of Mononucleosis between CDH mothers (n = 87) and controls (n = 809). Of the 87 women who gave birth to a baby with CDH, 3 (3.45%) had a diagnosis of infectious mononucleosis compared to 60 (6.90%) out of 809 controls. Our results showed no difference between the two groups (OR = 0.48, 95% CI [0.0946 – 1.5298], p=0.2632).

**Figure 4:**
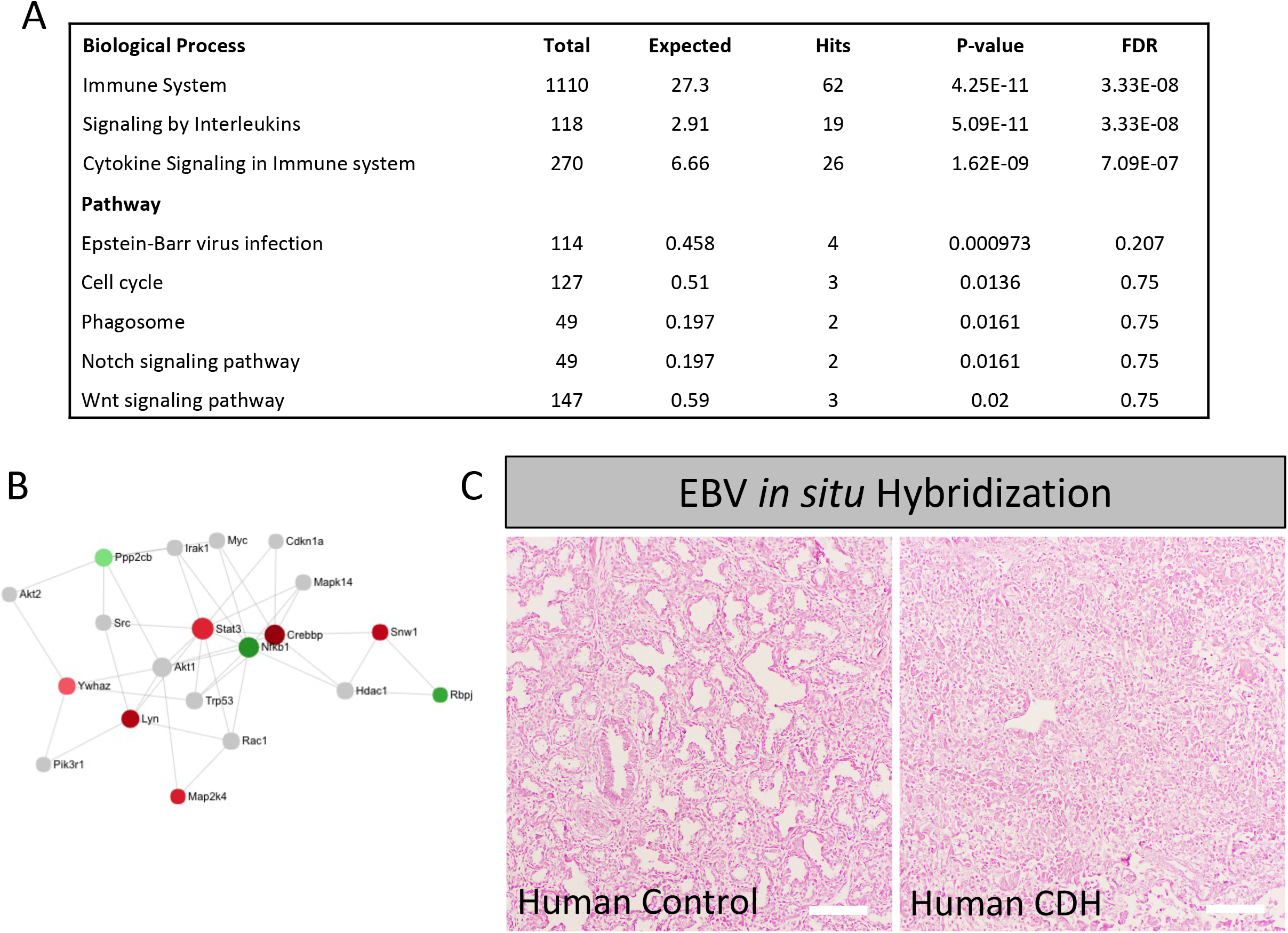
Pathway analysis determines an enrichment in processes associated to inflammation and immune response. A) Pathway analysis of significantly (Log2FC >1.5; p<0.05) altered proteins from the proteomics dataset of nitrofen (n=5) and control (n=5) lungs B) Protein network highlighting proteins that were significantly altered between nitrofen and control lungs and associated with Epstein Bar Virus C) Representative image showing *in-situ* hybridization against Epstein Bar Virus mRNA (EBER probe) in fetal lung tissues of fetopsy cases from CDH patients and age matched controls (n=10 CDH, n=5 controls). Scale bars 100μm.

### 3.4. STAT3 inhibition can rescue peripheral lung budding after direct nitrofen exposure ex vivo

As our results clearly demonstrated that STAT3 was increased in airway epithelium of nitrofen lungs (Figure 3D), we examined whether STAT3 inhibition could reverse the negative effects of direct nitrofen exposure on lung branching. Correia-Pintos’ group has previously shown that different STAT proteins are expressed during lung development and that STAT3 inhibition can enhance normal fetal lung growth *ex vivo* ^23^. Here, we tested whether STAT3 inhibition can partially rescue lung budding after *ex vivo* nitrofen exposure to E14.5 lung explants. In this *ex vivo* model we detected an increase of pSTAT3 protein in nitrofen exposed lungs after 2 and 3 days in culture (Figure 5A). Moreover, we detected a significant mRNA elevation of the known STAT3 downstream target, SOCS3. (Figure 5B). Nitrofen reduced peripheral lung buds and total lung size compared to control lung explants, whereas mesenchymal thickness was increased in nitrofen lungs (Figure 5C–F and Suppl. Figure 1). Adding STAT3 inhibitor (S3I-201) 6 hours after nitrofen treatment increased peripheral lung budding and restored the mesenchymal width back to the level of unexposed lungs (Figure 5C–F). In contrast, the total lung size was decreased after STAT3 inhibition when compared to control and nitrofen lungs (Figure 5D). Finally, we determined that STAT3 inhibition decreased pSTAT3 protein in nitrofen lungs to control levels (Figure 5G).

**Figure 5:**
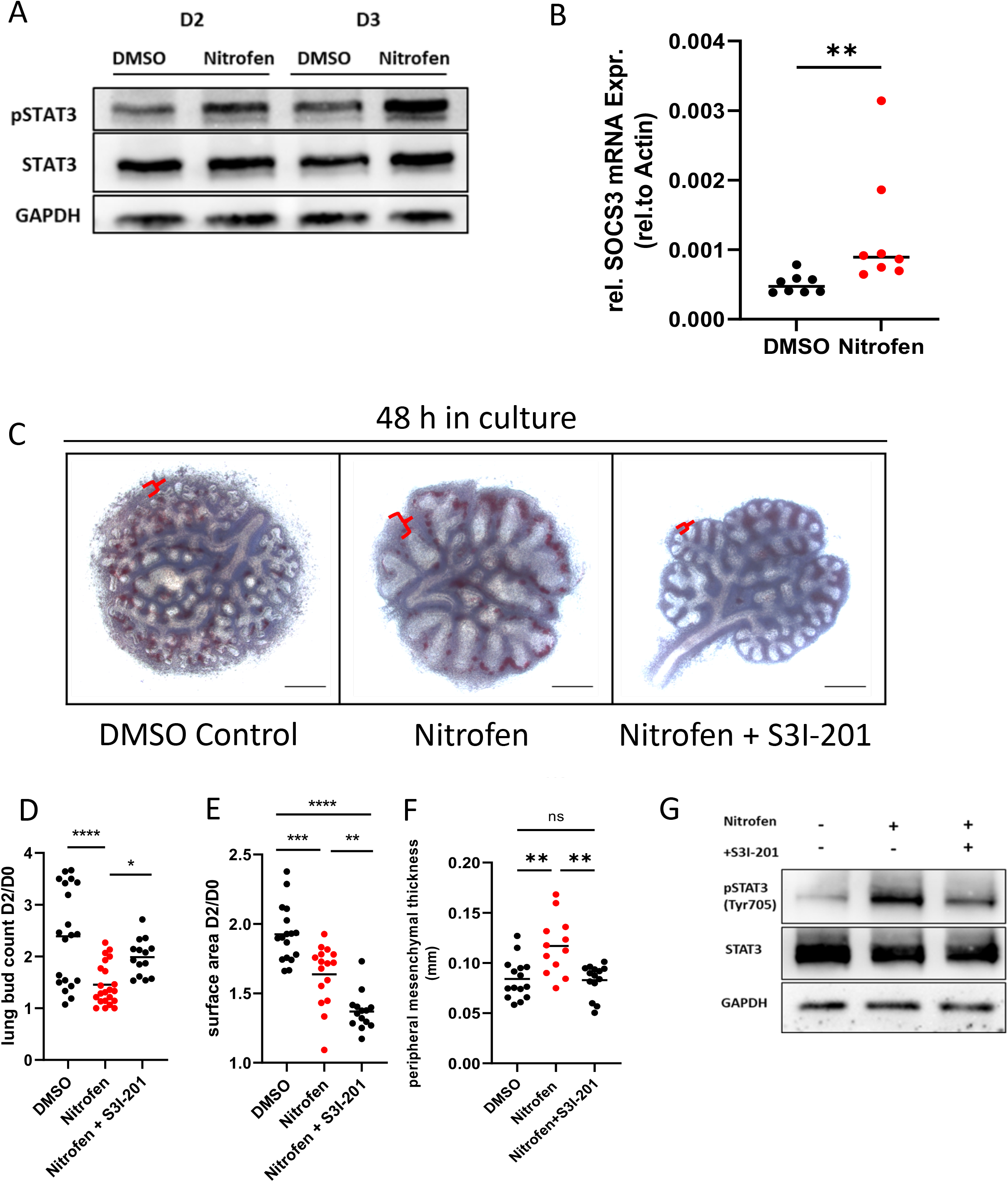
STAT3 inhibition can rescue peripheral lung budding after direct nitrofen exposure ex vivo. Fetal rat lungs were explanted at E14.5 and cultured for 48 – 72 hours in control medium, DMSO (vehicle), nitrofen medium or nitrofen medium + STAT3 inhibitor. A) Western blot for STAT3 and pSTAT3 in lung explants after 48h and 72h in culture. B) Relative mRNA expression of SOCS3, a downstream target of STAT3 in control and nitrofen lung explants after 48h. C) Representative images of lung explants after 48h in culture. Red brackets highlighting mesenchymal thickness D) Quantification of lung bud count as a ratio of number of peripheral lung buds after 48h in culture in relation to the number of lung buds at 0h in culture (n>25 lungs per group). E) Quantification of total lung size as a ratio of lung size after 48h in culture in relation to the lung size at 0h in culture (n>25 lungs per group) F) Quantification of mesenchymal thickness in mm after 48 h in culture. G) Representative Western blot for STAT3 and pSTAT3 in control, nitrofen and nitrofen + STAT inhibitor after 48 hours in culture. ANOVA test was used for statistical analysis, * p<0.05, ** p<0.01, *** p<0.001, ****p<0.0001. Scale bars 500μm.

### 3.5 Inflammatory cytokine profiles in tracheal fluids of CDH survivors are different compared to that in non-survivors

Informatics analysis of our proteomics data (described above) indicated an enrichment in pathways associated with cytokine signaling in nitrofen-induced hypoplastic lungs in the CDH animal model. It is known, that different cytokines are involved in lung developmental processes and regulation of surfactant production in type II pneumocytes ^24–26^. To test whether the cytokine milieu is changed between patients that survive CDH (n=4) and non-survivors (n=4), we examined a total of 42 cytokines in fetal tracheal fluid samples from CDH patients undergoing FETO at 28 weeks’ gestation (Suppl. Table 1). We detected an increase of G-CSF (p < 0.05), IL-15 (p < 0.05), IL-9 (p < 0.01) and IL-2 (p < 0.05) in fetal tracheal aspirates of CDH survivors compared to non-survivors (Figure 6A). We created a correlation heat map to determine correlation patterns between all 42 analysed cytokines. We observed an increased number of overall positive correlations between the individual cytokines in survivors (Figure 6B). Correlations between individual cytokines were altered between groups, e.g., in survivors EGF, RANTES and IL-1α were negatively correlated with the rest of the cytokines, while FGF2 and IL-R1 were negatively correlated to the other analytes in non-survivors. These data suggest that changes in specific cytokine abundance, as well as relative ratios of different cytokines, in tracheal aspirates can be correlated to specific CDH outcomes such as survivors vs non-survivors.

**Figure 6:**
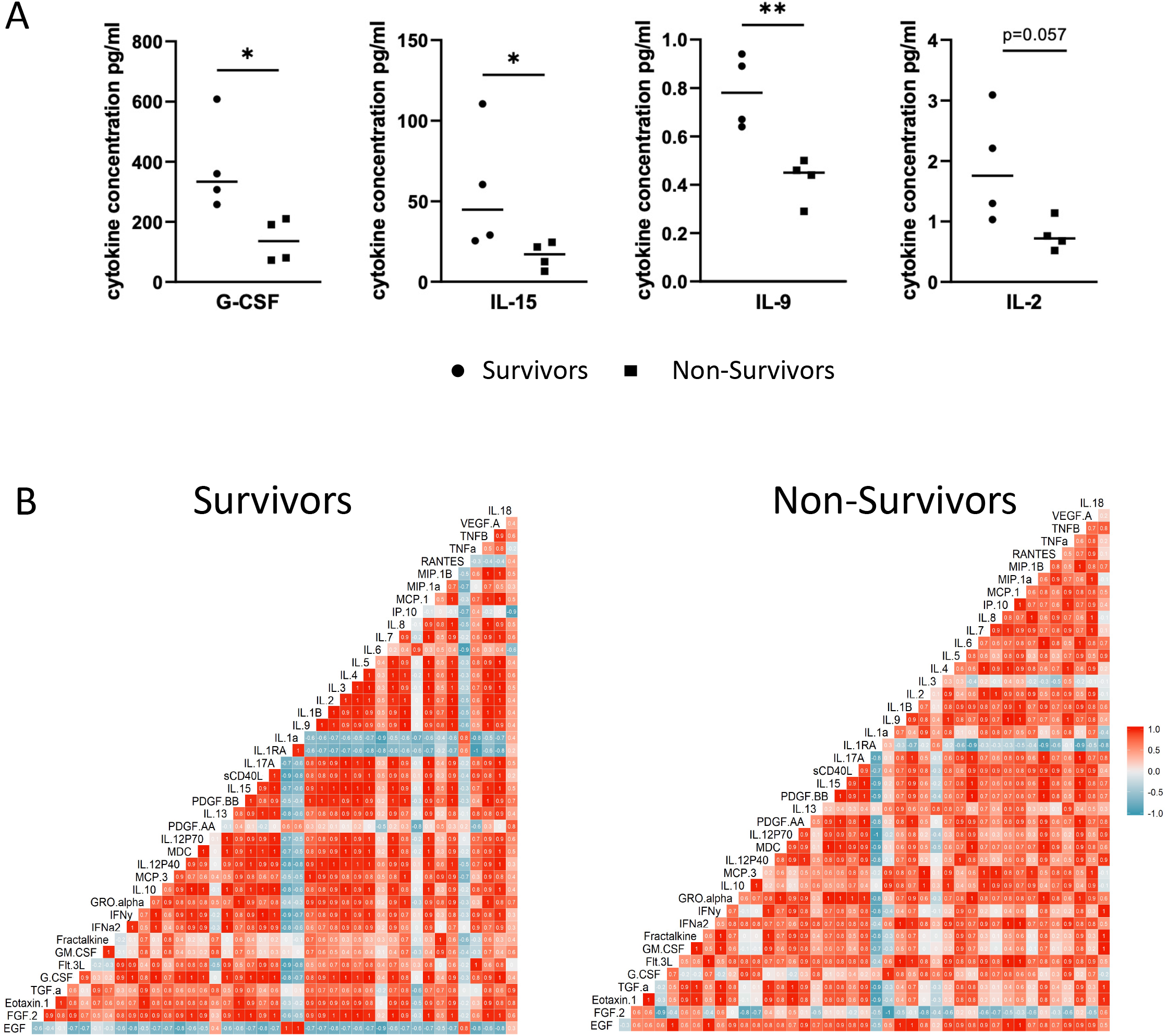
Inflammatory cytokine profiles in tracheal fluids of CDH survivors are different compared to that in non-survivors. A) Relative abundance of G-CSF, IL-15, IL-2, and IL-9 in tracheal fluids from CDH survivors and non-survivors that were obtained during occlusion of the fetal trachea via FETO. n=4 per group B) Correlation Heatmap showing the correlation between individual cytokines measured by 42 plex cytokine array using Pearson correlation. Squares represent individual correlation of two cytokines in all samples of each group. Red = positive correlation, blue = negative correlation.

## Discussion

Despite the phenotypic similarities between the nitrofen model and human CDH, the molecular underpinnings of the underlying abnormal lung development are insufficiently understood. Here we provide new insights into pathobiological processes that are altered in CDH. Using a quantitative proteomics approach, we found that inflammatory pathways and antimicrobial response elements are amongst the most prominent biological processes altered in nitrofen CDH lungs. This signature reflects alterations of key inflammatory molecules STAT3, CREBBP, NF-B LYN, and Tenascin C. Previously, Fleck et al. demonstrated that pro-inflammatory markers such as IL-1, IL-6 and IFN-α were significantly increased in cord blood of CDH neonates ^27^. Others showed that differentially expressed circulating microRNAs in CDH patients can be linked to inflammatory pathways and STAT3 signalling ^28^. Moreover, prenatal treatment with glucocorticoids led to a significant improvement of lung morphology in nitrofen CDH, indicating that anti-inflammatory strategies can be beneficial in experimental CDH ^29^.

Our proteomic analysis revealed a significant enrichment of virus associated pathways in nitrofen induced lung hypoplasia. We speculated that nitrofen exposure during early development can trigger pathways in rat fetal lungs that are similar to an innate immune response to viruses in human CDH. Some viruses can cross the placenta and cause different malformations in the fetus (i.e. human cytomegalovirus, Zika virus) ^30,31^. Although there is up to 100% of seroprevalence of human cytomegalovirus in the general population, related birth defects occur only in rare cases, suggesting that additional factors contribute to virus related congenital malformations ^30,32^. Although we cannot provide direct evidence to link CDH pathogenesis to a viral stimulus and associated innate immune response during lung development, we propose this as a potential mechanism that requires further investigation.

We report that STAT3 and pSTAT3(Tyr705) are significantly increased in nitrofen lungs corroborating previous studies in the nitrofen model showing increased STAT3 ^23,33^. Piairo et al. previously reported the enhancing effects of STAT3 inhibition to lung budding in normal fetal lung explants ^23^. We demonstrate that STAT3 inhibition can restore distal lung budding in nitrofen exposed hypoplastic lungs, associated with a decrease in mesenchymal thickening and thereby reduction of total lung size compared to nitrofen and control lungs. At E15, STAT3 is predominantly expressed in the mesenchyme of rat lungs ^23^. It is thus tempting to speculate, that abnormally enhanced STAT3 in nitrofen exposed lungs contributes to mesenchymal hyperproliferation and compromised cellular differentiation, a key feature of hypoplastic CDH lungs.

We also profiled cytokines in fetal tracheal aspirates from CDH patients undergoing fetoscopic tracheal occlusion (FETO) that eventually survived CDH compared to non-survivors. Thereby, we found changes in abundance of different cytokines and in cytokine correlation analysis between the groups, some of which are associated with STAT3 activation ^34–36^. Fetal tracheal aspirates can only be obtained in severe CDH cases undergoing this prenatal intervention in specialised centers ^37^, thus we are lacking a control group without CDH. However, our observations correspond to a study in a CDH sheep model reporting that cytokine and inflammatory response pathways were prominently enriched in tracheal fluids of CDH fetuses ^38^. We detected a significant increase of IL-2, IL-9, IL-15 and G-CSF in tracheal fluids of CDH survivors compared to non-survivors, suggesting a potential as biomarkers for disease severity and prognostication in cases undergoing FETO.

Using an unbiased proteomics approach and robust data analysis, we determined decreased Tenascin C as the most discriminant protein between hypoplastic CDH and control lungs. Tenascin C can promote branching morphogenesis and vascularization and regulate epithelial to mesenchymal transition, a process that is associated with CDH lung hypoplasia ^39^. We show that Tenascin C is significantly suppressed in nitrofen CDH lungs, which is in line with findings from a rabbit model of CDH ^40^. We were able to confirm decreased Tenascin C abundance in human fetal CDH lung samples, further demonstrating the validity of the animal model. Tenascin C is known to be expressed during embryonic development, and can regulate inflammatory responses engaging STAT3 and NF-κB pathways ^41^. These findings suggest that a decrease in Tenascin C may be one of the mechanisms associated with an increase in inflammatory responses in CDH lungs, engaging STAT3 as a central signaling intermediate in this process.

Several limitations of our study have to be acknowledged. Although we provide evidence for significant protein alterations in nitrofen CDH lungs, we are still unable to perform similar studies using human samples due to technical limitations when using fixed tissues. The translatability of findings from the nitrofen model to human CDH is restricted and results have to be interpreted with caution. Moreover, our study cannot fully address whether the proteomic signature seen in our model reflects an intrinsic lung defect in CDH or results from external compression to the developing lung. Future studies will have to investigate which cell type drives this signature associated with the lung phenotype in human CDH and if modulation of identified pathways can rescue abnormal lung development *in vivo*. Overall, this study provides a comprehensive proteomic analysis of experimental CDH lungs, identifies proteins and signaling intermediates associated with the underlying processes, and describes the interface between inflammation and CDH. We also demonstrate that intervening in a key inflammatory signaling intermediate STAT3, which is increased in CDH, can be beneficial in hypoplastic lungs.

## Acknowledgements

The authors gratefully acknowledge Ying Lao for technical assistance with mass spectrometry and Victor Spicer at the Manitoba Centre for Proteomics and Systems Biology for the LC-MS/MS and related data analysis. We thank Jacquelyn Schwartz for their help with the animal experiments and immunohistochemistry. We also thank Micheline Philippot – Rosset for the EBV *in-situ* hybridization. Furthermore, we thank Dr. John Wilkins for intellectual input into the proteomics study design. We thank Mandy Laube for their suggestions for the lung explant culture system. We also thank the families of patients who consented for their participation and cooperation.

## Author contributions

RW, NM and RK conceived the study. RW, PL, AT, JR, NP, JD, MM, DP and CS obtained the samples for the study. RW and HP performed the LC/MS-MS sample preparation. RW, HP, PL, JR, AT and CP analyzed the data. HP and RW performed the validation experiments. PL and NP performed the *ex vivo* experiments. RW, PL, HP, AT and MM performed immunological staining experiments. GD and SLM performed the cohort study. RW, HP, NM and RK wrote the manuscript. RK was the senior lead for this study and extensively edited the manuscript together with RW, NM, CP and ML. ML, JG, JR, JD and CP provided substantial intellectual input for the study. All authors reviewed and edited the final manuscript.

---

**Supplemental Figure 1:**
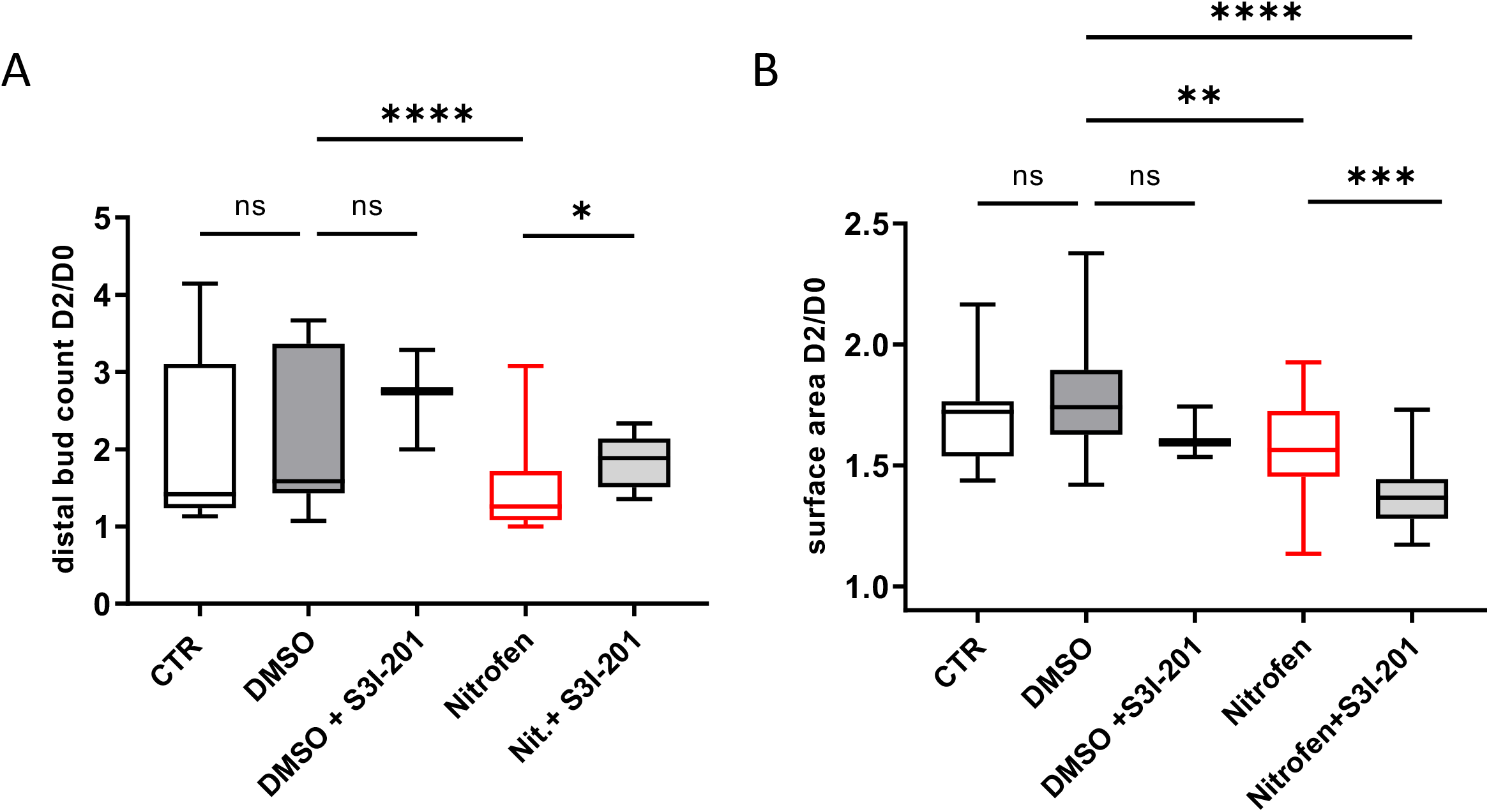
Lung explants with quantification of distal lung buds and lung surface area as a ratio of Day2/Day0 in culture. Controls (medium only), DMSO (vehicle only), DMSO + STAT3 inhibitor, nitrofen and nitrofen + STAT3 inhibitor. n>3 per group, ANOVA test was used for statistical analysis, * p<0.05, ** p<0.01, *** p<0.001, ****p<0.0001.

**Supplemental Table 1:** Demografic information of patients undergoing FETO.

|  | Survivors (n=4) | Non-Survivors (n=4) | P-value |
| --- | --- | --- | --- |
| Gestational Age at Plug (weeks) | 28.1 (28.7 - 27.1) | 27.9 (29.1 - 27.0) | 0.84 |
| **Observed/ Expected Lung to Head Ratio (%)** | **22.5 (23.6 - 17.5)** | **21.6 (24.0 - 15.9)** | **0.87** |
| Liver herniated | 100% | 100% | 1.00 |
| Birth weight (g) | 2780 (3180 - 2160) | 3195 (3278 - 2650) | 0.3 |
| Day of neonatal death | N/A | 1 (2-0) |  |

